## Supplementary Table S2 for "Peptidoglycan-tethered and free forms of the Braun lipoprotein are in dynamic equilibrium in *Escherichia coli*"

**Table S2. Kinetic analysis of the content of Tri➔KR isotopologues in new and old Tri and KR moieties**

| Combinations of Tri➔KR isotopologues | Biological repeat | Relative abundance (%) at 0, 10, 20, 40, and 60 min: WT | | | | | Δ*yafK* | | | | |
|---|---|---|---|---|---|---|---|---|---|---|---|
| | | 0 | 10 | 20 | 40 | 60 | 0 | 10 | 20 | 40 | 60 |
| Containing light KR new➔new + old➔new | 1 | 0 | 6.57 | 14.68 | 31.38 | 47.85 | 0 | 6.17 | 15.74 | 38.71 | 58.45 |
| | 2 | 0 | 5.23 | 12.15 | 33.03 | 50.53 | 0 | 3.57 | 8.92 | 25.92 | 46.25 |
| | 3 | 0 | 3.21 | 10.39 | 43.32 | 53.95 | 0 | 3.46 | 10.38 | 26.81 | 55.20 |
| | 4 | | 4.95 | | | | | | | 29.48 | |
| | 5 | | 3.73 | | | | | | | | |
| | **Mean** | 0 | 4.74 | 12.41 | 35.91 | 50.77 | 0 | 4.40 | 11.68 | 29.93 | 53.30 |
| | **SD** | 0 | 1.61 | 2.25 | 10.19 | 4.36 | 0 | 2.17 | 3.97 | 6.93 | 7.23 |
| Containing heavy KR new➔old + old➔old | 1 | 100 | 93.43 | 85.32 | 68.62 | 52.15 | 100 | 93.83 | 84.26 | 61.29 | 41.55 |
| | 2 | 100 | 94.77 | 87.85 | 66.97 | 49.47 | 100 | 96.39 | 89.47 | 73.89 | 53.75 |
| | 3 | 100 | 96.79 | 89.61 | 56.68 | 46.05 | 100 | 96.65 | 89.53 | 76.52 | 44.80 |
| | 4 | | 95.05 | | | | | | | 70.52 | |
| | 5 | | 96.27 | | | | | | | | |
| | **Mean** | 100 | 95.26 | 87.59 | 64.09 | 49.23 | 100 | 95.62 | 87.75 | 70.56 | 46.70 |
| | **SD** | 0 | 3.06 | 6.14 | 7.60 | 3.30 | 0 | 1.80 | 5.30 | 8.49 | 9.05 |
| Containing light Tri new➔new + new➔old | 1 | 0 | 3.45 | 13.25 | 42.86 | 70.45 | 0 | 1.14 | 5.42 | 25.18 | 49.60 |
| | 2 | 0 | 4.03 | 12.25 | 49.73 | 73.44 | 0 | 1.55 | 1.95 | 14.97 | 37.91 |
| | 3 | 0 | 1.42 | 7.95 | 53.05 | 75.84 | 0 | 1.44 | 5.52 | 20.42 | 49.05 |
| | 4 | | 2.60 | | | | | | | 19.79 | |
| | 5 | | 1.31 | | | | | | | | |
| | **Mean** | 0 | 2.56 | 11.15 | 48.54 | 73.25 | 0 | 1.38 | 4.30 | 19.79 | 45.52 |
| | **SD** | 0 | 1.29 | 2.82 | 12.59 | 4.86 | 0 | 0.50 | 2.04 | 6.37 | 6.66 |
| Containing heavy Tri old➔new + old➔old | 1 | 100 | 96.55 | 86.75 | 57.14 | 29.55 | 100 | 98.86 | 94.58 | 74.82 | 50.40 |
| | 2 | 100 | 95.97 | 87.75 | 50.27 | 26.56 | 100 | 98.41 | 96.44 | 84.84 | 62.09 |
| | 3 | 100 | 98.58 | 92.05 | 46.95 | 24.16 | 100 | 98.67 | 94.39 | 82.92 | 50.95 |
| | 4 | | 97.40 | | | | | | | 80.21 | |
| | 5 | | 98.69 | | | | | | | | |
| | **Mean** | 100 | 97.44 | 88.85 | 51.46 | 26.75 | 100 | 98.65 | 95.13 | 80.70 | 54.48 |
| | **SD** | 0 | 3.38 | 5.58 | 5.21 | 2.80 | 0 | 3.47 | 7.22 | 9.05 | 9.63 |

1
