## Supplementary Table S1 for "Peptidoglycan-tethered and free forms of the Braun lipoprotein are in dynamic equilibrium in *Escherichia coli*"

**Table S1. Kinetic analysis of the relative abundance of Tri→KR isotopologues in wild-type and Δ*yafK***

| Tri→KR isotopologues | Biological repeat | Relative abundance (%) of isotopologues at 0, 10, 20, 40, and 60 min | | | | | | | | | |
|---|---|---|---|---|---|---|---|---|---|---|---|
| | | WT | | | | | Δ*yafK* | | | | |
| | | **0** | **10** | **20** | **40** | **60** | **0** | **10** | **20** | **40** | **60** |
| new→new | 1 | 0 | 0.57 | 3.26 | 17.72 | 38.11 | 0 | 0.40 | 2.52 | 17.26 | 39.85 |
| | 2 | 0 | 1.26 | 2.94 | 21.68 | 40.93 | 0 | 1.09 | 1.49 | 9.04 | 30.84 |
| | 3 | 0 | 0.42 | 1.80 | 33.55 | 45.44 | 0 | 0.72 | 2.76 | 10.21 | 40.14 |
| | 4 | | 0.47 | | | | | | | 13.37 | |
| | 5 | | 0.17 | | | | | | | | |
| | **Mean** | **0** | **0.58** | **2.67** | **24.32** | **41.49** | **0** | **0.74** | **2.26** | **12.17** | **36.94** |
| | **SD** | **0** | **0.41** | **0.77** | **8.24** | **3.69** | **0** | **0.34** | **0.68** | **4.45** | **5.29** |
| new→old | 1 | 0 | 2.88 | 9.99 | 25.14 | 32.34 | 0 | 0.74 | 2.90 | 7.92 | 9.75 |
| | 2 | 0 | 2.77 | 9.31 | 28.05 | 32.51 | 0 | 0.46 | 0.46 | 5.93 | 7.07 |
| | 3 | 0 | 1.00 | 6.15 | 19.50 | 30.41 | 0 | 0.72 | 2.76 | 10.21 | 8.91 |
| | 4 | | 2.12 | | | | | | | 6.42 | |
| | 5 | | 1.14 | | | | | | | | |
| | **Mean** | **0** | **1.98** | **8.48** | **24.23** | **31.75** | **0** | **0.64** | **2.04** | **7.62** | **8.58** |
| | **SD** | **0** | **0.88** | **2.05** | **4.35** | **1.17** | **0** | **0.15** | **1.37** | **1.92** | **1.37** |
| old→new | 1 | 0 | 6.00 | 11.42 | 13.65 | 9.73 | 0 | 5.77 | 13.22 | 21.45 | 18.60 |
| | 2 | 0 | 3.97 | 9.21 | 11.36 | 9.60 | 0 | 2.48 | 7.43 | 16.88 | 15.41 |
| | 3 | 0 | 2.79 | 8.59 | 9.77 | 8.51 | 0 | 2.74 | 7.62 | 16.61 | 15.06 |
| | 4 | | 4.48 | | | | | | | 16.11 | |
| | 5 | | 3.56 | | | | | | | | |
| | **Mean** | **0** | **4.16** | **9.74** | **11.59** | **9.28** | **0** | **3.66** | **9.42** | **17.76** | **16.36** |
| | **SD** | **0** | **1.20** | **1.49** | **1.95** | **0.67** | **0** | **1.83** | **3.29** | **2.48** | **1.95** |
| old→old | 1 | 100 | 90.55 | 75.33 | 43.49 | 19.81 | 100 | 93.09 | 81.36 | 53.37 | 31.80 |
| | 2 | 100 | 92.01 | 78.54 | 38.92 | 16.97 | 100 | 95.93 | 89.01 | 67.96 | 46.67 |
| | 3 | 100 | 95.79 | 83.45 | 37.19 | 15.64 | 100 | 95.93 | 86.77 | 66.31 | 35.89 |
| | 4 | | 92.93 | | | | | | | 64.10 | |
| | 5 | | 95.13 | | | | | | | | |
| | **Mean** | **100** | **93.28** | **79.11** | **39.86** | **17.47** | **100** | **94.98** | **85.71** | **62.94** | **38.12** |
| | **SD** | **0** | **2.18** | **4.09** | **3.25** | **2.13** | **0** | **1.64** | **3.93** | **6.57** | **7.68** |
