## Supplementary File 1 for "Peptidoglycan-tethered and free forms of the Braun lipoprotein are in dynamic equilibrium in *Escherichia coli*"

**BW25113**
**t = 0 min**

Tri→KR
$m/z$ = 607.347
$z$ = 2

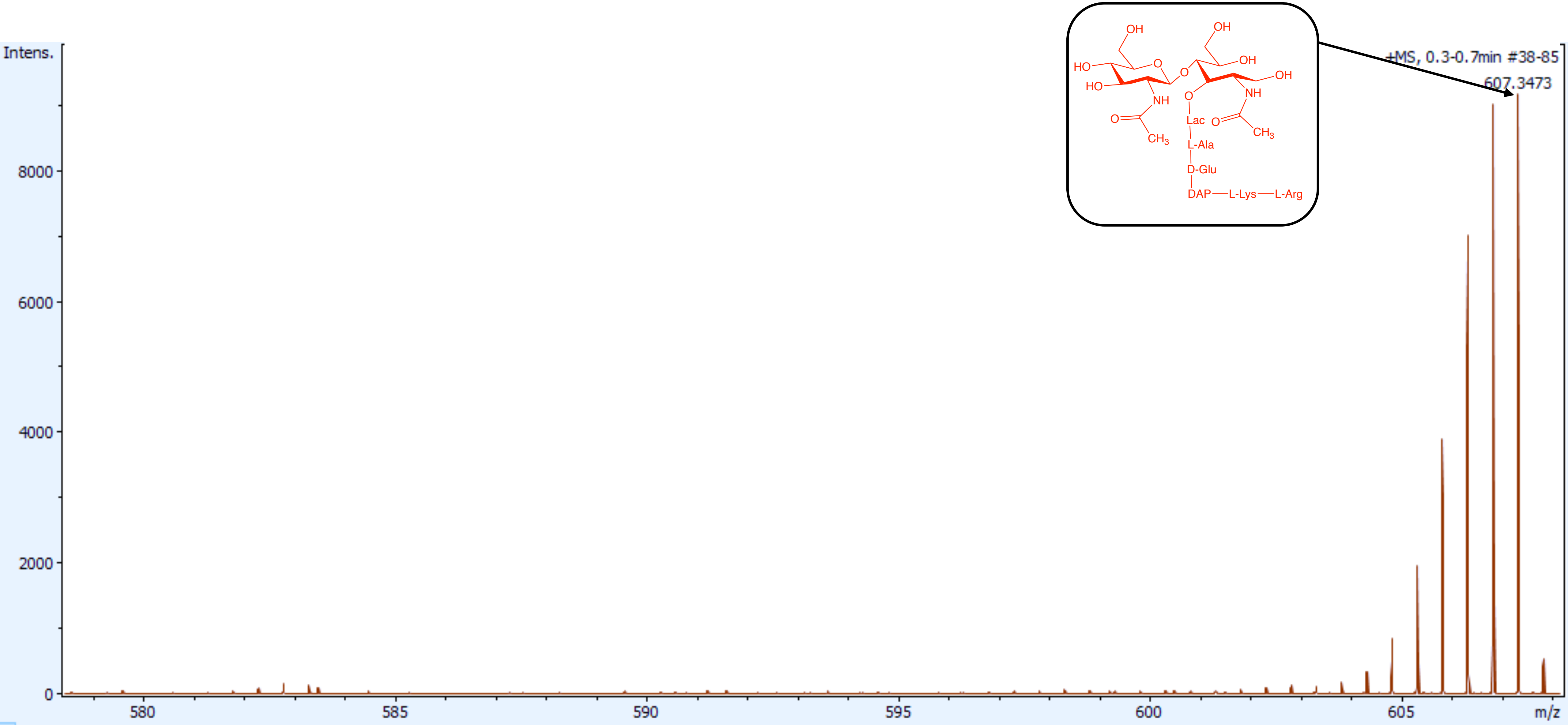

**BW25113**
**t =5 min**

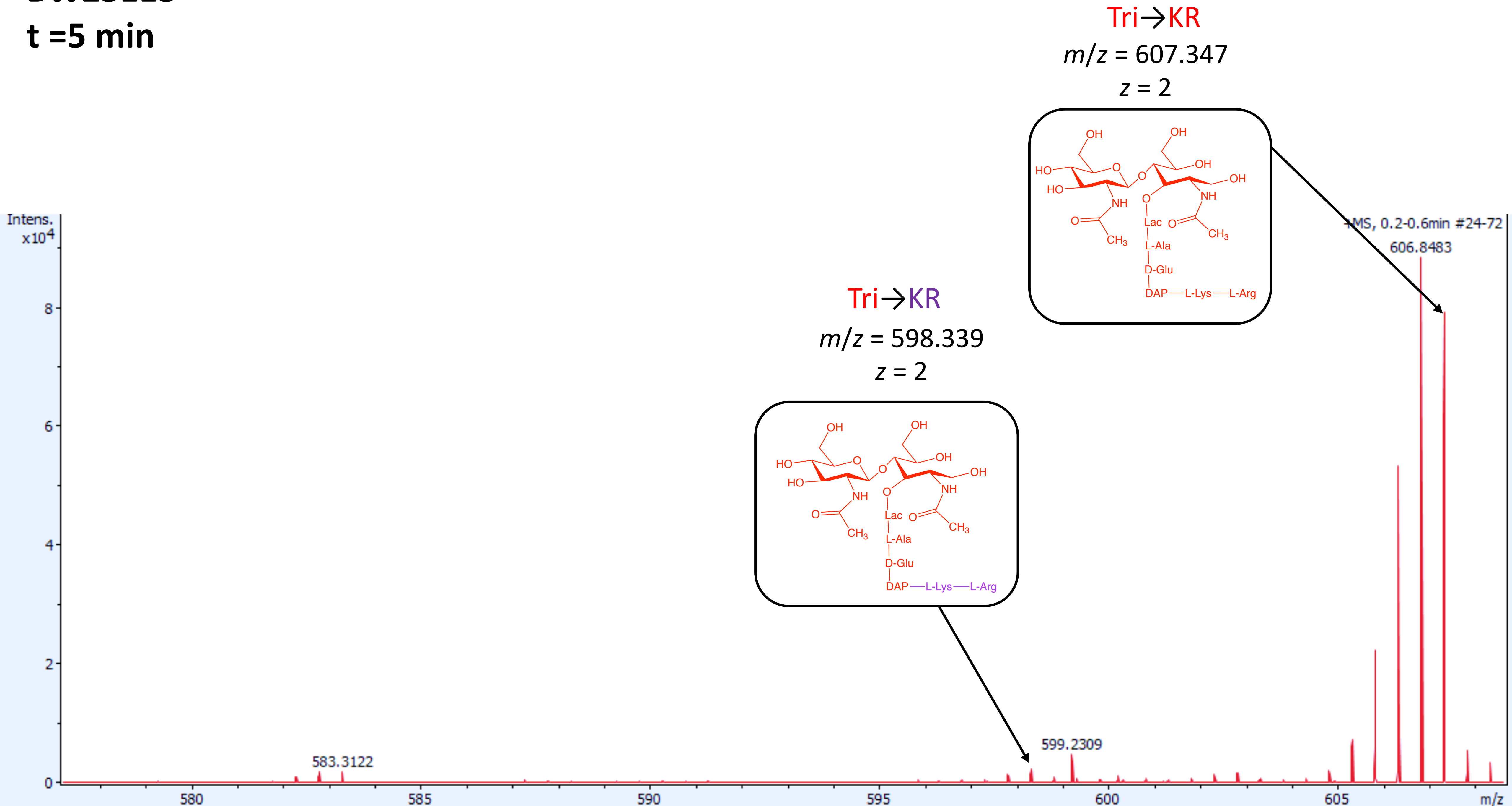

**BW25113**
**t = 10 min**

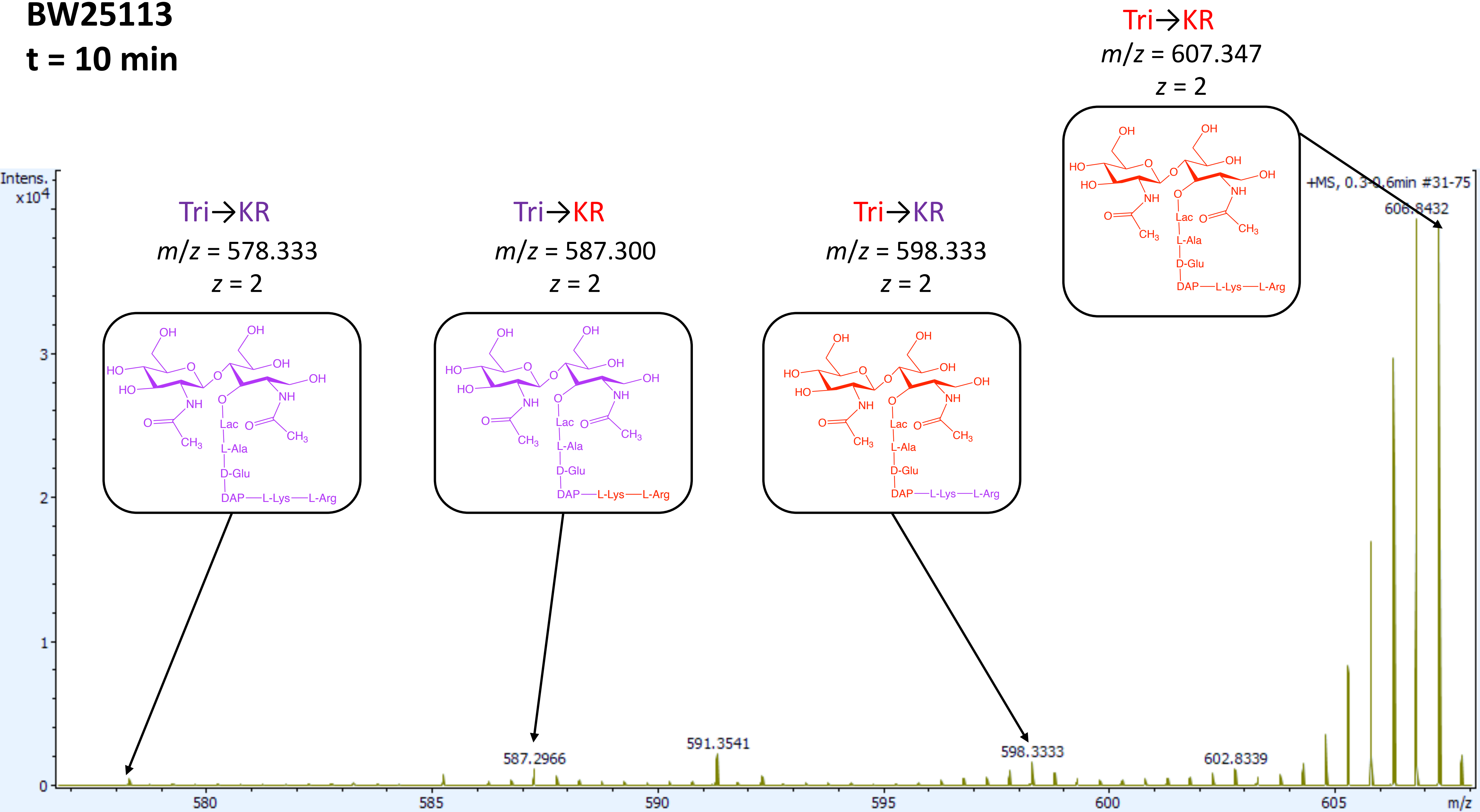

# BW25113
### t = 20 min

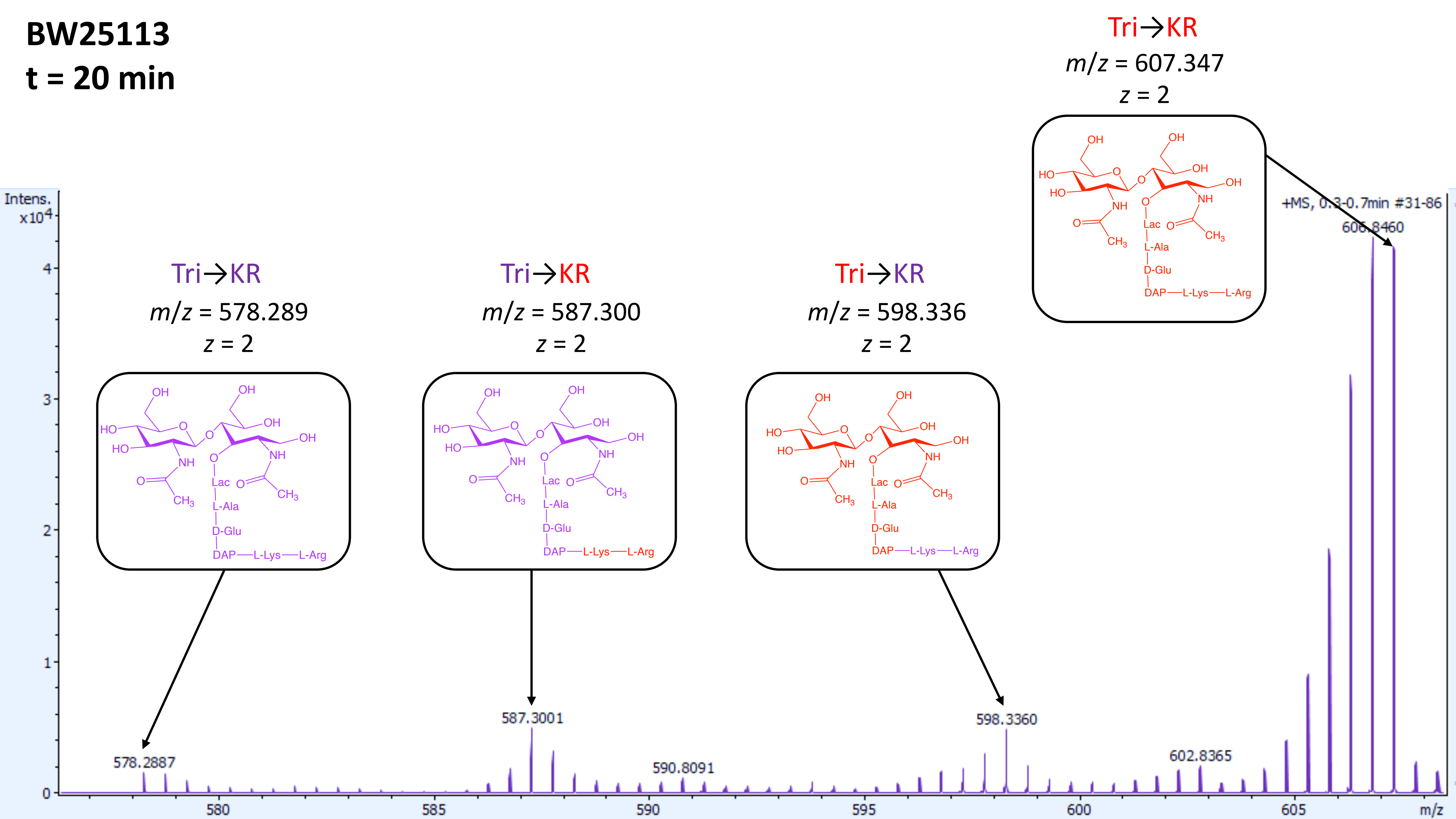

**BW25113**

**t = 40 min**

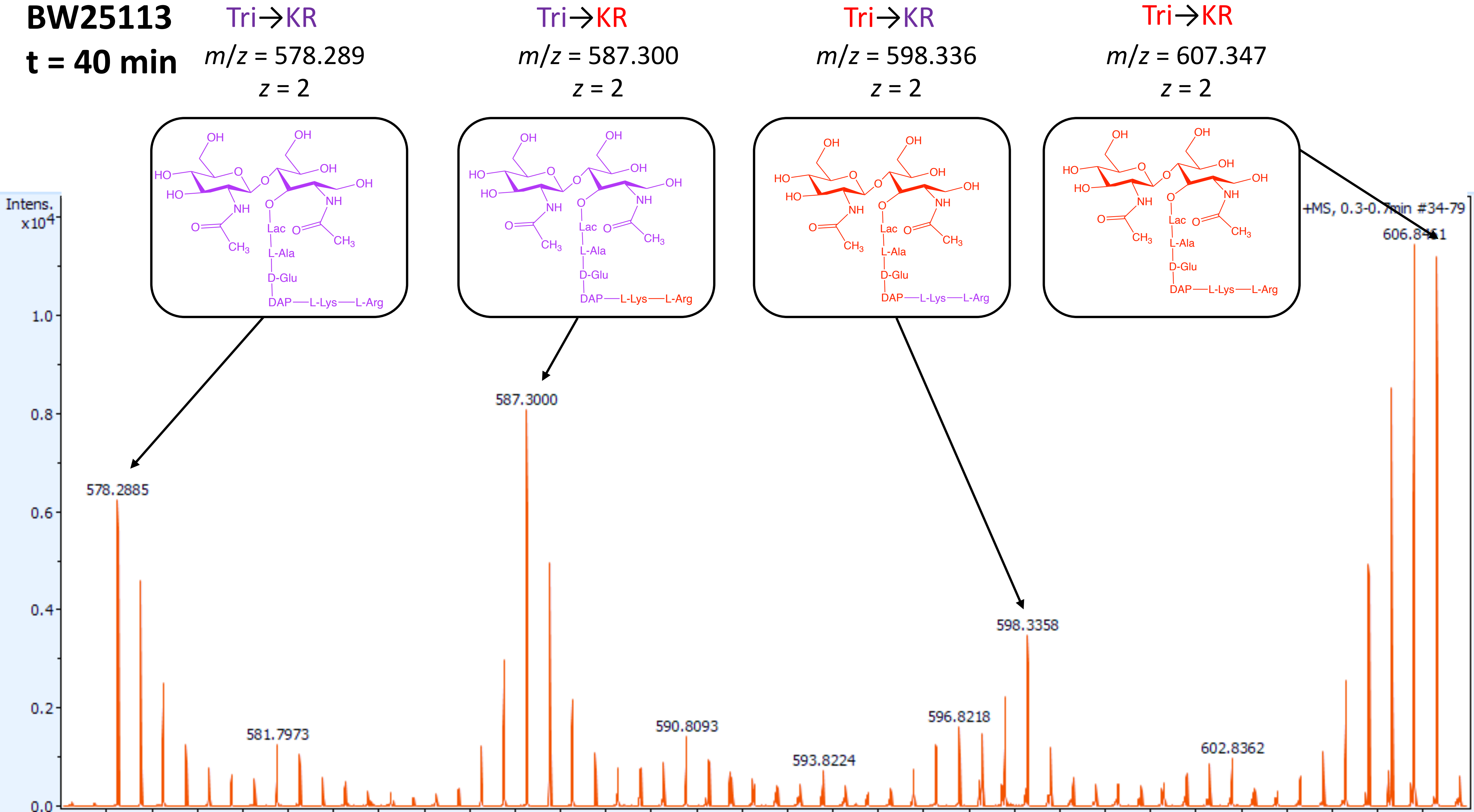

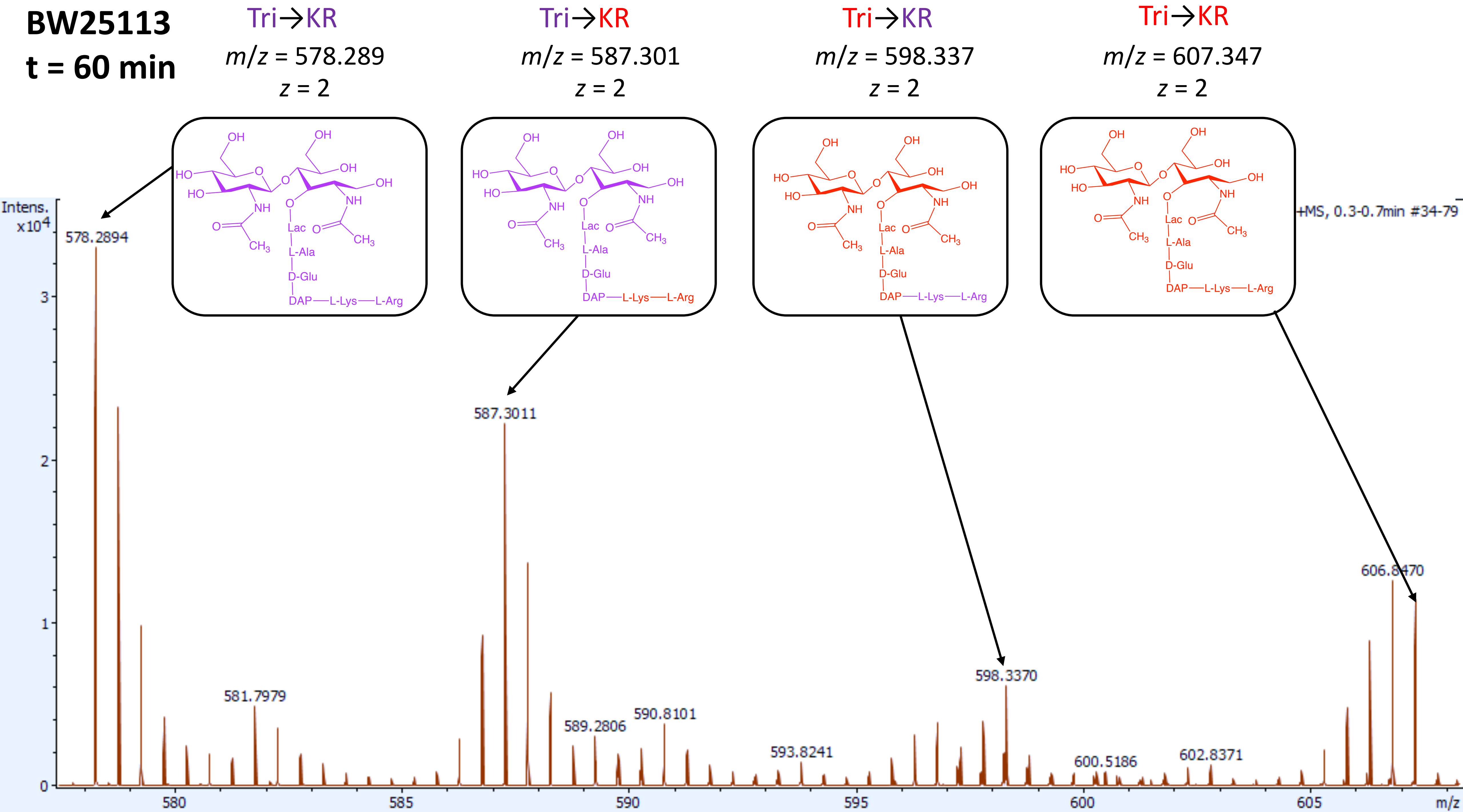

BW25113
t = 60 min
Tri→KR
m/z = 578.289
z = 2
Tri→KR
m/z = 587.301
z = 2
Tri→KR
m/z = 598.337
z = 2
Tri→KR
m/z = 607.347
z = 2
Lac
L-Ala
D-Glu
DAP—L-Lys—L-Arg
Intens.
x10^4
+MS, 0.3-0.7min #34-79
578.2894
581.7979
587.3011
589.2806
590.8101
593.8241
598.3370
600.5186
602.8371
606.8470
580
585
590
595
600
605
m/z

***ΔyafK***

**t = 0 min**

Tri→KR

$m/z$ = 607.347

$z$ = 2

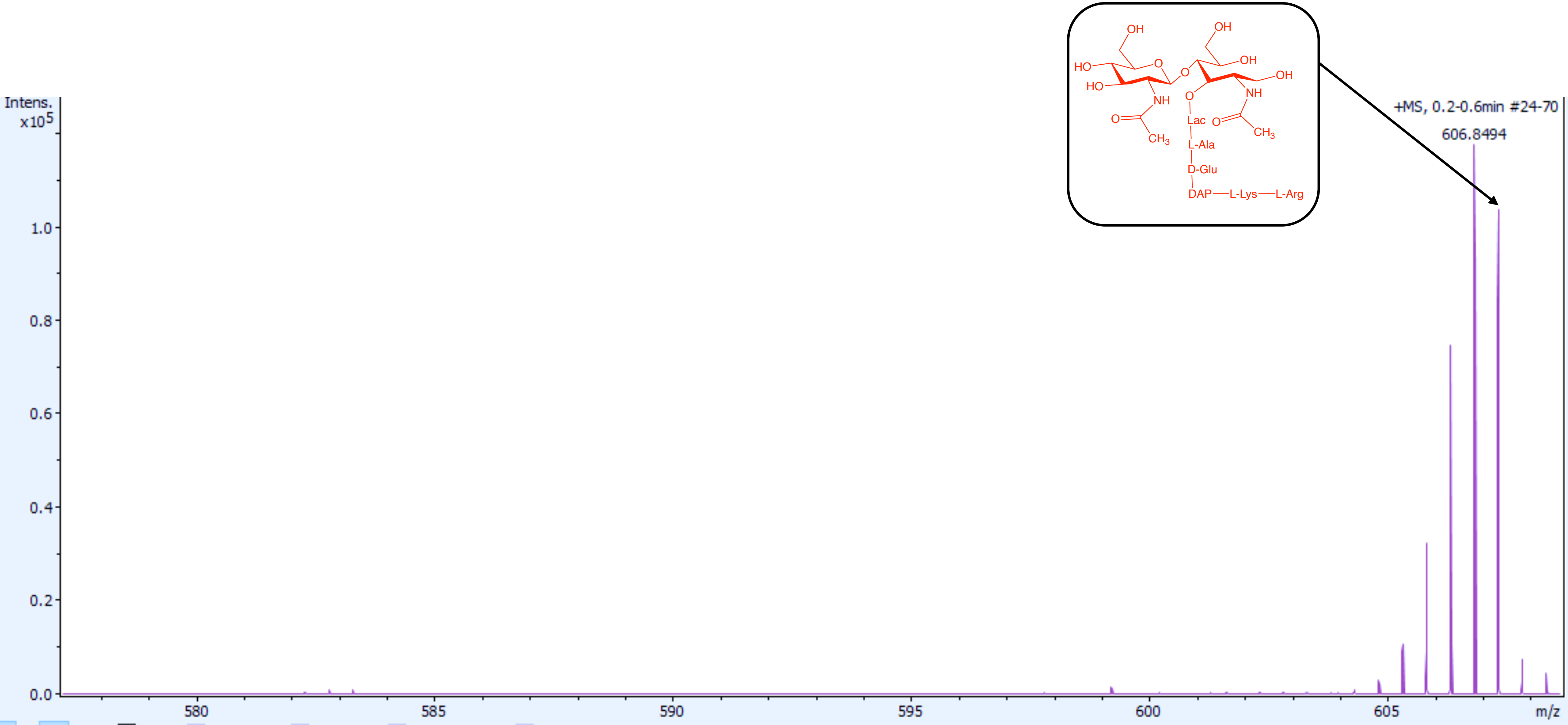

***ΔyafK***
**t = 5 min**

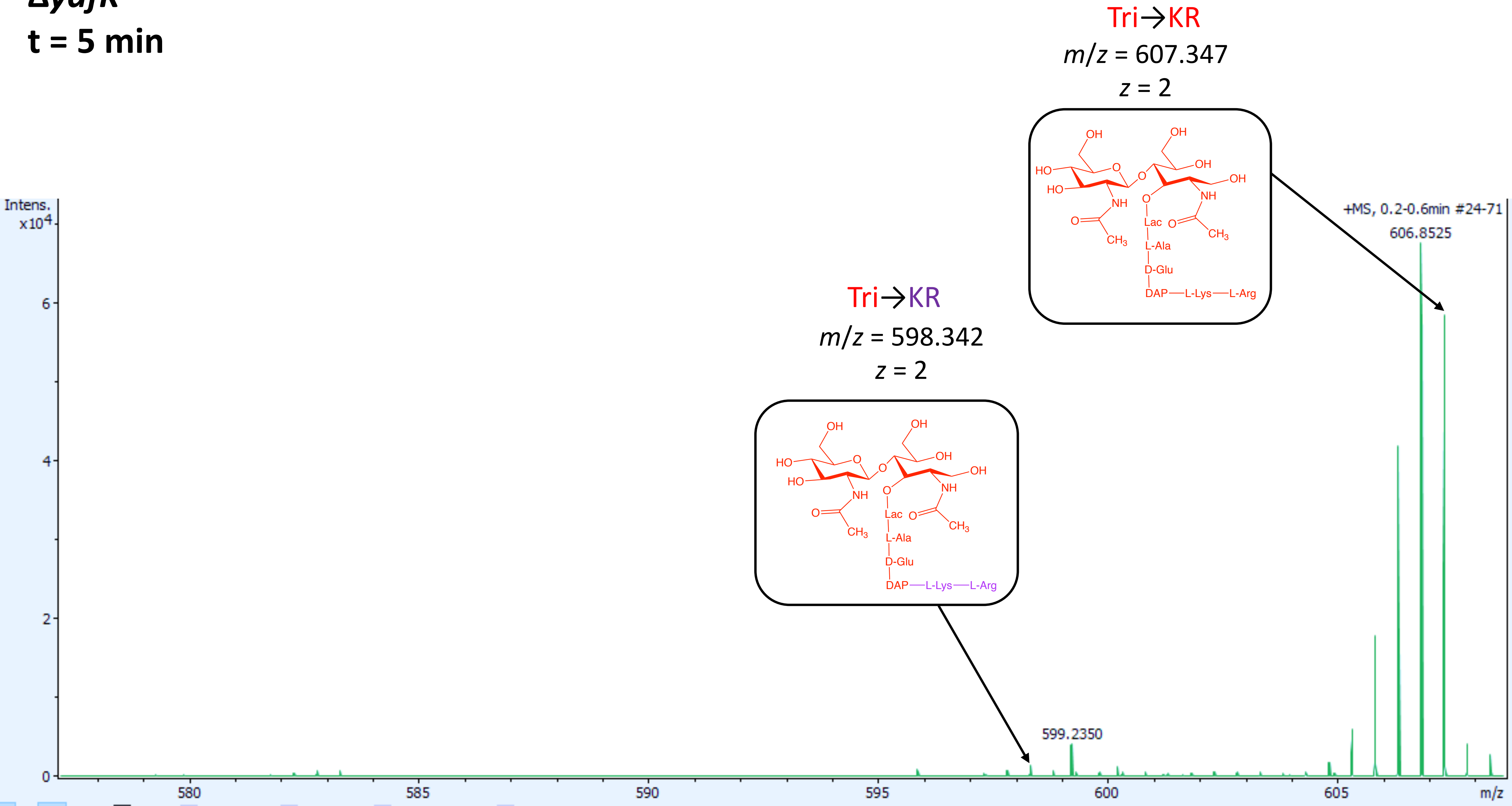
Tri→KR
m/z = 607.347
z = 2
OH
OH
HO
O
O
OH
OH
HO
NH
O
NH
O
Lac
O
CH3
CH3
L-Ala
D-Glu
DAP—L-Lys—L-Arg
Tri→KR
m/z = 598.342
z = 2
DAP—L-Lys—L-Arg
Intens.
x10^4
+MS, 0.2-0.6min #24-71
606.8525
599.2350
6
4
2
0
580
585
590
595
600
605
m/z

***ΔyafK***
**t = 10 min**

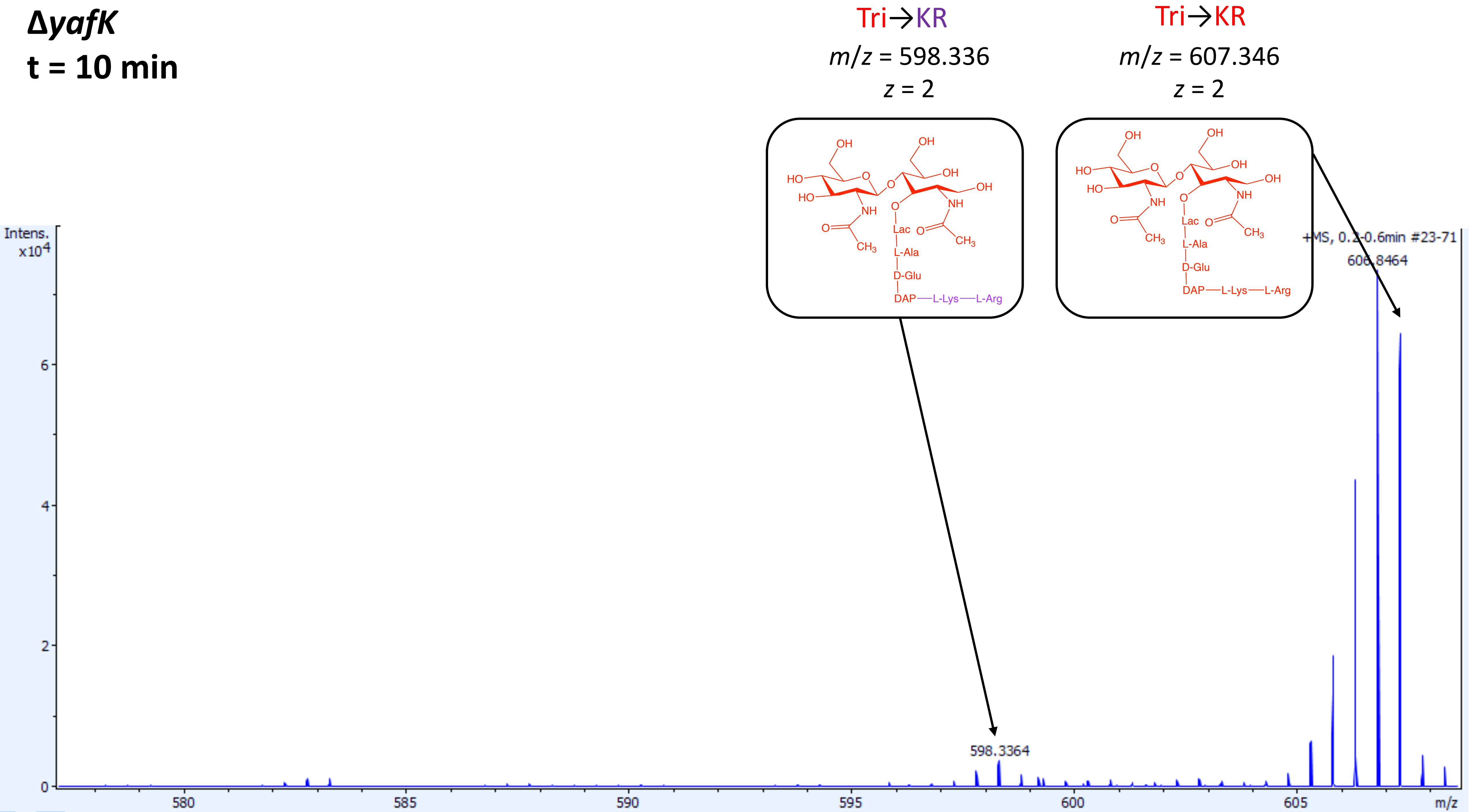

**Δ*yafK***
**t = 20 min**

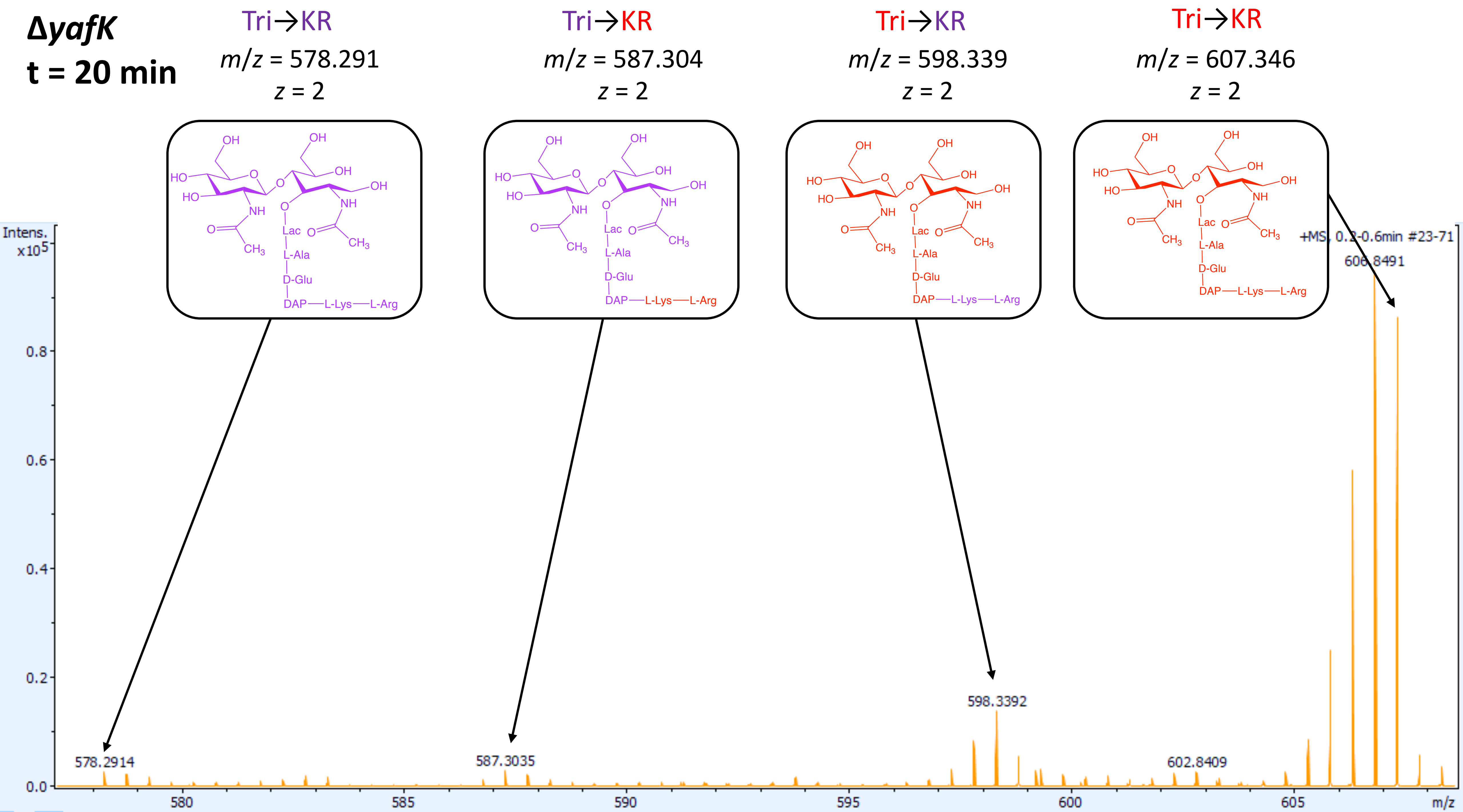

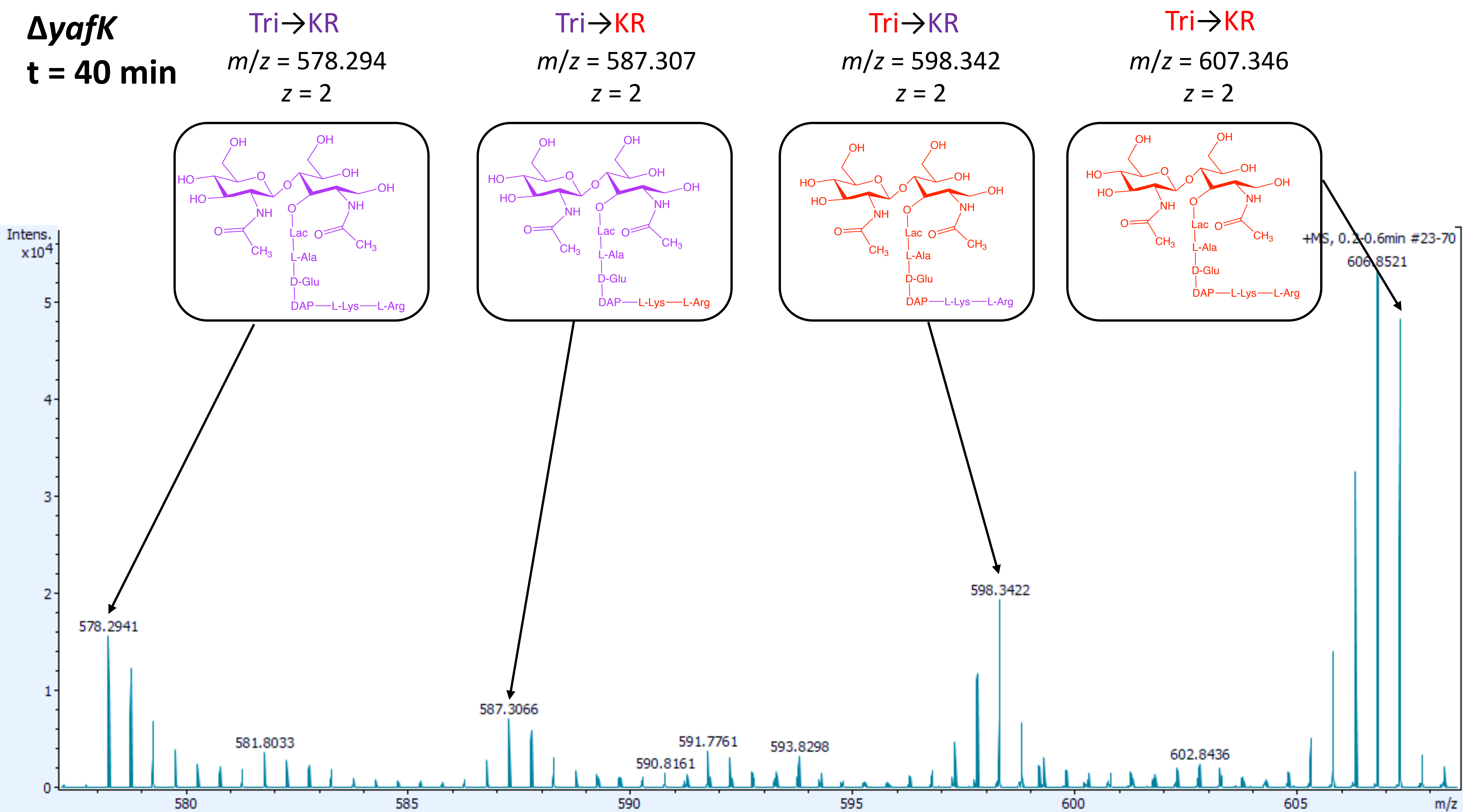

ΔyafK
t = 40 min
Tri→KR
m/z = 578.294
z = 2
Tri→KR
m/z = 587.307
z = 2
Tri→KR
m/z = 598.342
z = 2
Tri→KR
m/z = 607.346
z = 2
Lac
L-Ala
D-Glu
DAP—L-Lys—L-Arg
Intens.
x10^4
+MS, 0.2-0.6min #23-70
578.2941
581.8033
587.3066
590.8161
591.7761
593.8298
598.3422
602.8436
606.8521
m/z

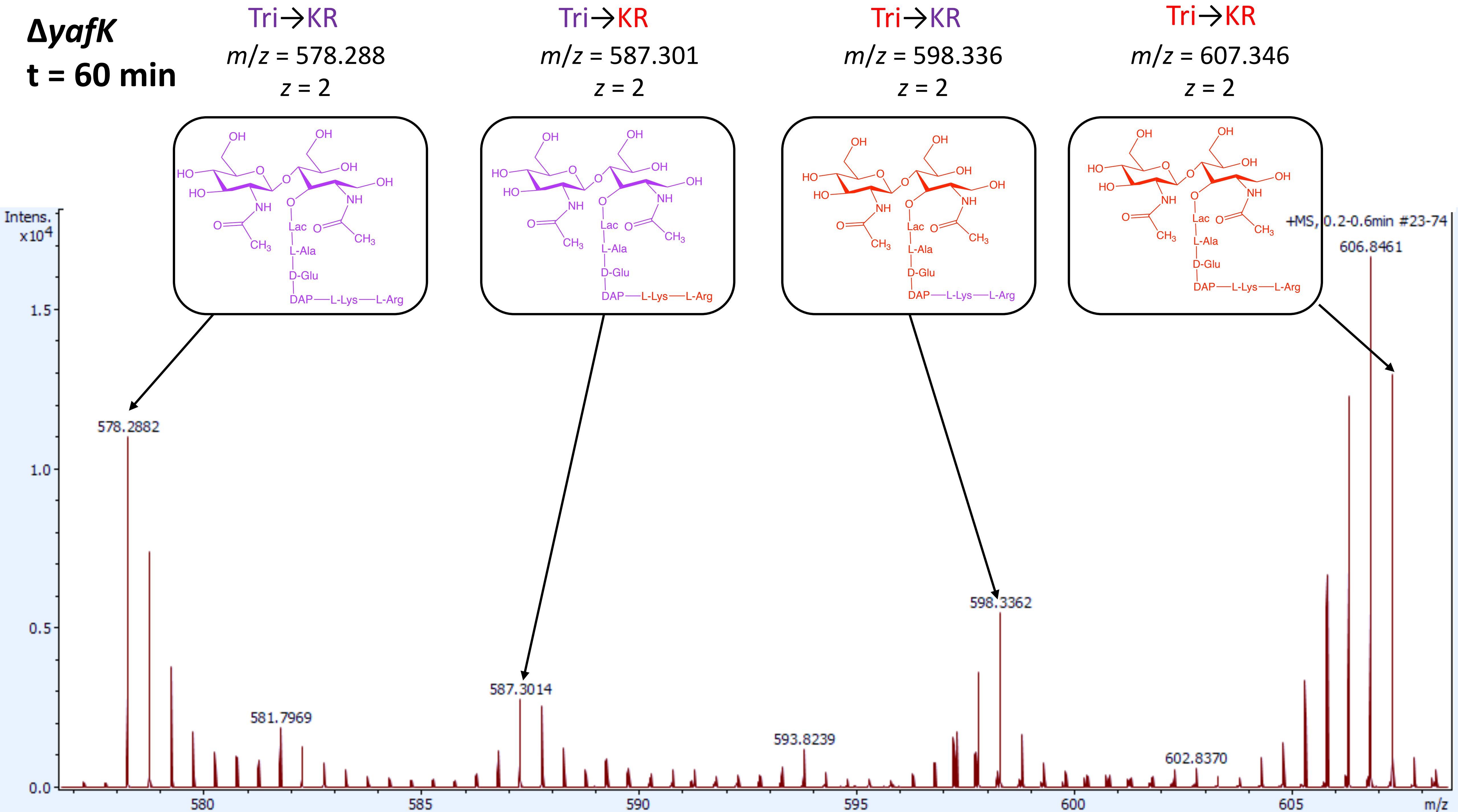
ΔyafK
t = 60 min
Tri→KR
m/z = 578.288
z = 2
Tri→KR
m/z = 587.301
z = 2
Tri→KR
m/z = 598.336
z = 2
Tri→KR
m/z = 607.346
z = 2
Lac
L-Ala
D-Glu
DAP—L-Lys—L-Arg
Intens.
x10^4
+MS, 0.2-0.6min #23-74
606.8461
578.2882
581.7969
587.3014
593.8239
598.3362
602.8370
m/z
